# S1P induces bleb-based T cell motility via S1PR1-dependent activation of RhoA and WNK1

**DOI:** 10.1101/2025.10.03.680076

**Authors:** Franklin Staback Rodriguez, Henry De Belly, Yichen Zhang, Evelyn Strickland, Gordon L. Frazer, Judith Ojukwu, Benjamin J. Williamson, Orion D. Weiner, Janis K. Burkhardt

## Abstract

In vivo, the chemokine CCL19 and its receptor CCR7 control T cell retention in lymph nodes, while the lipid chemoattractant spingosine-1-phosphate (S1P) drives T cell egress from lymphoid organs. CCL19 is known to activate actin polymerization at the leading edge of migrating cells, generating a mode of motility driven by lamellipodial protrusions. In contrast, we showed recently that S1P induces a transient lamellipodial response, followed by pressure-driven bleb-based motility. Here, we elucidate the mechanisms controlling S1P responses in naïve T cells. We show that S1P signals through S1PR1, with coupling through Gai. In contrast to CCR7, which signals through Gai to induce sustained Rac1 activation, S1PR1 engagement yields only weak and transient Rac1 activation; the dominant response is sustained activation of RhoA. This pathway, together with a pathway involving phospholipase C and myosin light chain kinase, results in phosphorylation of myosin regulatory light chain (MLC) and enhanced myosin contractility. Inhibition of mTORC2 blocks MLC phosphorylation, consistent with evidence that tension sensing by mTORC2 can couple Rac1 and RhoA signaling during leukocyte migration. Surprisingly, although RhoA pathway inhibitors blocked S1P-induced MLC phosphorylation and blebbing, they failed to block S1P-dependent chemotaxis. This led to the identification of a second arm of the S1P response: WNK1-dependent phosphorylation of SPAK1 and OSXR1, proteins that regulate ion channels and water influx. Partial WNK1 inhibition, together with inhibition of myosin contractility, was sufficient to block S1P-induced blebbing and chemotaxis, indicating that S1P-driven T cell migration involves coordinate activation of myosin contractility and water influx.

**One sentence summary:** S1P signals elicit sustained RhoA activation and water influx to drive bleb-based T cell motility.\

## Introduction

To carry out their role in immune defense, T cells must move throughout the body in a highly choreographed fashion (*1, 2*). From the bloodstream, naïve T cells enter secondary lymphoid organs where they migrate through the stroma in search of dendritic cells presenting cognate antigen. T cells that fail to find cognate antigens within a lymphoid organ egress and re-enter another lymphoid organ (directly or via the circulatory system) to continue surveillance. In contrast, T cells that detect antigen proliferate and differentiate into effector cells. These effector cells also egress from lymphoid organs, eventually re-entering circulation. Subsequently, they extravasate at sites of inflammation to enter the tissues and fight infection (*3*). These trafficking patterns are directed by a large group of chemoattractants, which signal through G-protein coupled receptors (GPCRs) to induce the cytoskeletal changes needed for directed T cell motility (*4, 5*). One of the best studied chemoattractants is the chemokine CCL19, which signals through its receptor CCR7 to direct T cell movement within the stroma of lymphoid organs (*6, 7*). This action is opposed by the lipid chemoattractant sphingosine-1-phosphate (S1P), which induces T cell egress (*6, 8*). On naïve T cells, the primary receptor for S1P is S1PR1 (*9, 10*). S1P engagement of S1PR1 induces receptor endocytosis, desensitizing the cells to further signaling as they travel the body (*11, 12*). Once T cells return to lymphoid tissues, the low local concentration of S1P allows receptor re-expression and another round of egress and circulation. T cell trafficking in response to S1P has been intensively studied at the organismal level, but much less is known about the cell biological mechanisms through which S1P controls T cell motility.

Our lab recently discovered that CCL19 and S1P induce fundamentally distinct modes of T cell motility (*13*). Migration induced by CCL19 and other conventional chemokines is powered by actin-polymerization and leading-edge lamellipodial protrusions (*14*). This mode of motility, which is also observed in T cells migrating randomly on ICAM-1, allows T cells to migrate long distance on 2D surfaces even under shear flow conditions and permits passage across vascular endothelial barriers (*15–17*). In contrast, T cells stimulated with S1P exhibit an alternate mode of migration. Initially, S1P-stimulated cells form lamellipodial protrusions similar to those induced by CCL19. However, within about 3 minutes of S1P stimulation, T cells begin to generate leading edge blebs, which drive forward movement. Motile responses to S1P are short-lived (*13*), and it has been suggested that S1P drives short-range chemokinesis rather than chemotaxis (*18*). This could suffice to allow egress from lymphoid organs.

The signaling pathways through which CCL19 controls lamellipodial motility are well studied. The CCL19 receptor, CCR7, signals through Gai, inducing activation of Rac1 and its effector Wave2, which then drives Arp2/3-dependent branched-actin polymerization (*19, 20*). These events generate membrane ruffles and leading edge lamellipodial protrusions.

Leukocytes undergoing lamellipodial motility also exhibit low levels of active Rho, which induces myosin II contractility in the trailing uropod (*21, 22*). In contrast with CCL19, little is known about the molecular mechanisms underlying T cell responses to S1P. S1PR1 is the dominant S1P receptor in naïve T cells, and our previous work showed that this receptor is important for migration in our system (*13*). In addition, we showed S1P induces myosin-light chain (MLC) phosphorylation, resulting in myosin contractility and elevated intracellular pressure (*13*).

However, the signaling pathways linking these events to one another were unexplored. In other cell types, similar myosin-dependent blebbing responses occur downstream of RhoA and phospholipase C (PLC) (*23–25*) . However, S1PR1 is coupled to Gai, which typically activates Rac1 rather than RhoA (*26, 27*). Therefore, we sought to understand how signaling through S1PR1 could elicit myosin activation in naïve T cells, and to directly test the functional consequences of these events for T cell migration.

In the current study, we compare the signaling events induced by CCL19 and S1P in naïve T cells, and we delineate the cytoskeletal regulatory pathways through which S1P induces bleb-based motility. We find that whereas CCL19 signals through CCR7 and Gai to induce sustained Rac1 activation, S1P signals through S1PR1 and Gαi to induce transient Rac1 activation followed by sustained RhoA activity. In keeping with this, we find that the S1P-dependent increases in myosin contractility are due to activation of RhoA signaling, together with PLC. In parallel with this, S1PR1 engagement activates a WNK1 signaling pathway that promotes water influx and works together with myosin contractility to induce bleb formation. This study represents a comprehensive analysis of the signaling pathways through which S1P drives naïve T cell migration, and it reveals that S1P and CCL19 initiate fundamentally distinct cytoskeletal signaling cascades, leading to complementary motile mechanisms.

## Results

### S1P induces naïve T cell chemotaxis

We showed previously that stimulation of naïve murine T cells with S1P induces a polarized blebbing response associated with myosin contractility and elevated intracellular pressure (*13*). Since the S1P response is much shorter-lived than responses directed by CCL19, we proposed that the mode of motility induced by S1P might allow T cells to migrate through short constrictions such as those found at egress sites from lymphoid organs (*13*). Subsequent work by Garcia-Seyda et al (*18*) using microfluidic systems led to similar conclusions for human T cells. Due to the short-lived S1P polarity responses and the short distances the cells traversed, those authors argued that S1P induces chemokinesis rather than chemotaxis. However, even short-range migratory responses can be directional, and such directionality might be very important for efficient egress responses. We therefore conducted transwell assays to address this issue. Naïve CD4^+^ T cells were isolated from mice and cultured overnight in charcoal-stripped FBS, which resensitizes the cells to S1P signaling (*13*). Cells were then added to the top wells of transwell chambers and allowed to settle on the filters. S1P was then added to the bottom wells, to the top wells along with cells, or to both wells of the chamber. Transmigration occurred only when S1P was added only to the bottom well, consistent with a true chemotactic response (Fig. 1A).

**Figure 1:**
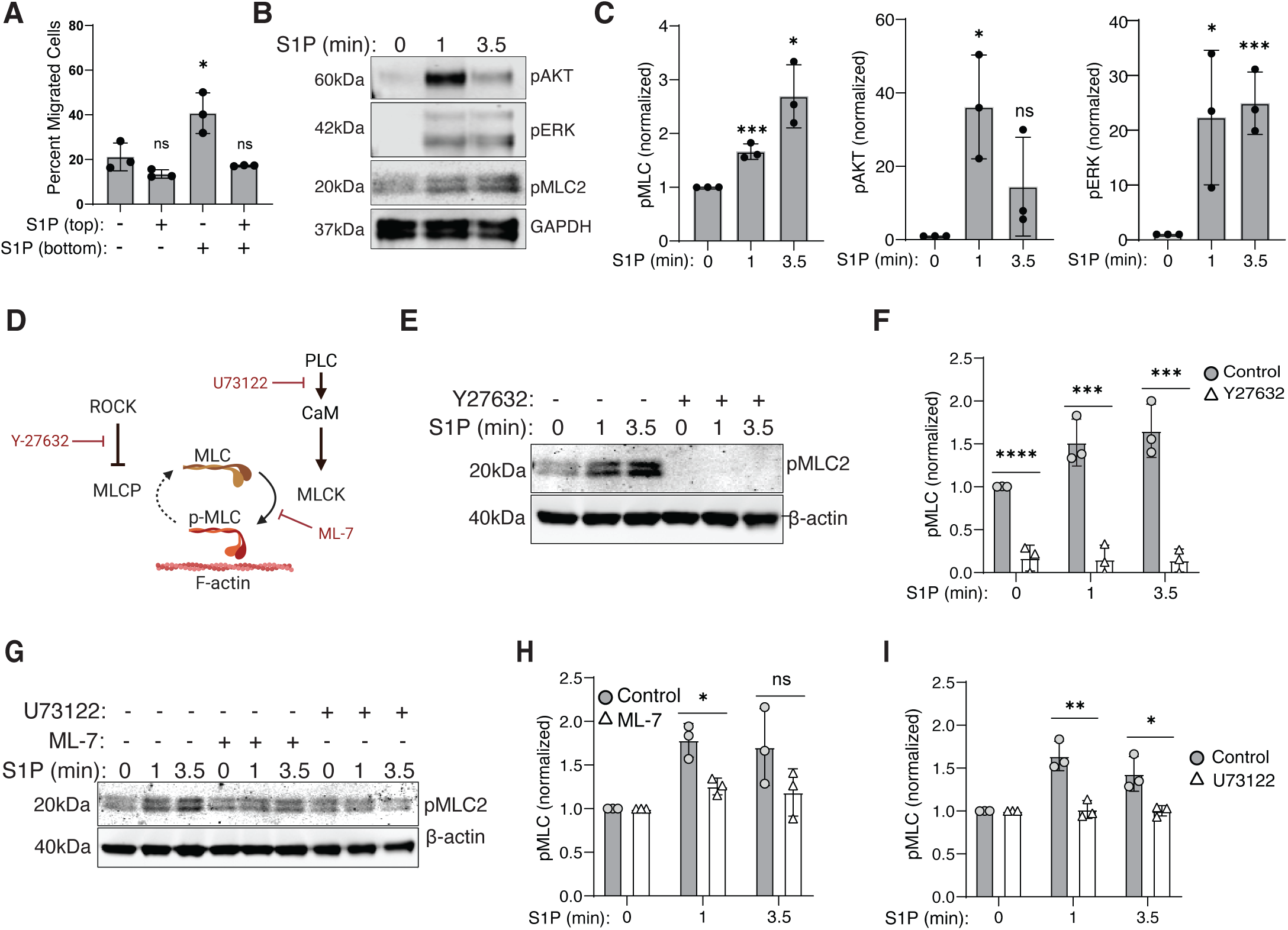
S1P-dependent myosin contractility is controlled by ROCK and PLC. (A) T cells chemotax toward S1P. Naïve CD4+ T cells were sensitized by overnight culture in CS-FBS, and chemotactic responses to 10nM S1P were measured using transwell assays. Significant migration was only observed when S1P was present only in the bottom well. n=3 biological replicates. (B-C) S1P activates phosphorylation of Akt, Erk, and MLC. (B) Naïve CD4+ T cells were stimulated with S1P for the indicated times, lysed with hot SDS, and analyzed by immunoblotting for the indicated phospho-proteins. GAPDH was used as a loading control. (C) Quantification of multiple experiments performed as shown in panel B (n=3). Band intensities were quantified and normalized to GAPDH. Fold increases were determined relative to the unstimulated control for each experiment. (D) Canonical model for MLC phosphorylation. RhoA activates Rho kinase (ROCK), which phosphorylates and inhibits MLC phosphatase. Additionally, PLC-mediated Ca^2+^ release activates calmodulin, which activates MLC kinase to phosphorylate MLC. Increased pMLC levels activate actomyosin contractility. Inhibitors are indicated in magenta. (E-F) pMLC increase occurs downstream of ROCK. T cells were cultured for 20min with the ROCK inhibitor Y-27632 or control media prior to stimulation with S1P. Cells were lysed with hot SDS and immunoblotted for pMLC. β-actin was used as a loading control. (F) Quantification of experiments performed as in panel E. Fold comparisons were drawn to the unstimulated (untreated) control (n=3). (G-I) pMLC increase occurs downstream of PLC and MLCK. T cells were pretreated with PLC inhibitor U73122, the MLCK inhibitor ML-7, or control media, stimulated for the indicated times with S1P, and analyzed as in panel E. (H-I) Quantification of experiments performed as in panel G. Fold comparisons were drawn to the unstimulated controls for each treatment condition (n=3). For all panels, data represent means +/- StDev from three independent biological replicates. Statistical analysis was performed using an unpaired Student’s T test, relative to the unstimulated (for C) or untreated (for F,H,I) controls. *, p<0.05; **, p<0.01; ***, p<0.005; ****, p<0.001

### S1P signals through RhoA and PLC to activate myosin light chain phosphorylation

We next set out to define the signaling pathway through which S1P triggers this response. We began by confirming our previous findings that S1P treatment of naïve T cells induces phosphorylation of Akt, ERK, and myosin light chain (MLC) (*13*) . As we reported previously, S1P induced rapid phosphorylation of Akt at Ser473 and ERK1/2 at Thr202/Tyr204 (Fig. 1B and C). In addition, it induced phosphorylation of MLC at Thr18/Ser19, an event that is known to activate myosin contractility (*28, 29*). Since myosin function is needed for S1P-induced T cell blebbing (*13*), we next focused on elucidating the events that lead to MLC phosphorylation. MLC phosphorylation is canonically regulated by myosin light chain kinase (MLCK) and myosin light chain phosphatase (MLCP) (Fig. 1D). MLCP is negatively regulated by Rho kinase (ROCK), functioning downstream of RhoA. Additionally, ROCK can directly phosphorylate RhoA (*30*). To ask whether the RhoA/ROCK/MLCP pathway controls S1P-driven MLC phosphorylation, we pre-incubated T cells with the ROCK inhibitor Y27632 and evaluated pMLC levels at baseline and after S1P stimulation. ROCK inhibition completely abrogated the baseline pMLC signal and prevented S1P-dependent phosphorylation (Fig. 1E and F). Since MLCK is activated by PLC via Ca^2+^ release and Calmodulin (CaM), we conducted similar experiments in the presence of the PLC inhibitor U73122 and the MLCK inhibitor ML-7. While these inhibitors did not perturb baseline pMLC levels, both prevented S1P-induced increases in MLC phosphorylation (Fig. 1G-I). Taken together, these results show that S1P signals through both arms of the canonical MLC regulatory pathway.

### S1P preferentially activates RhoA over Rac1 in naïve T cells

We showed previously that S1P-dependent migration of naïve T cells is mediated by S1PR1 (*13*), which is typically thought to drive Rac1 activation (*31–33*). However, our new data demonstrating the involvement of ROCK and MLCP suggest that S1P activates the RhoA signaling pathway in these cells. We therefore investigated RhoA and Rac1 activation directly. Naive T cells were stimulated with S1P for 0.5, 1, or 5 min, lysed and analyzed using a modified pull-down approach. Since CCL19 is well known to induce Rac1 activation, stimulation with CCL19 was performed for comparison. As anticipated based on downstream signaling events, stimulation with S1P induced significant increases in RhoA-GTP levels (Fig. 2A, light bars).

**Figure 2:**
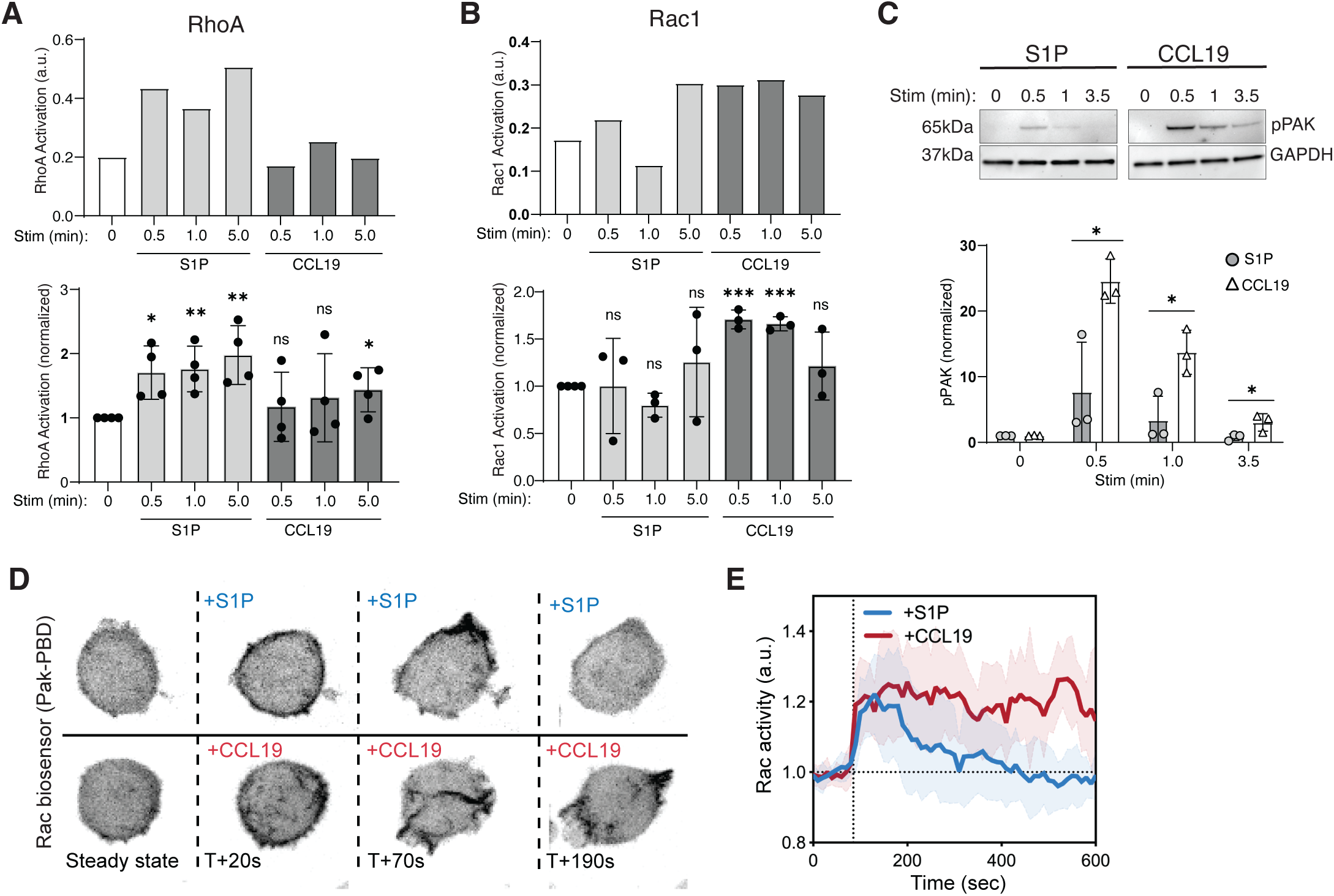
S1P induces strong and sustained RhoA activation and brief, low-level Rac1 activation. (A-B) Analysis of RhoA and Rac1 activation by S1P and CCL19. Naïve CD4+ T cells were stimulated with S1P or CCL19 for the indicated times, then rapidly lysed and analyzed for GTP-bound RhoA and Rac1. Top panels show representative experiments; bottom panels show averages of multiple biological replicates (n=4 for RhoA; n=3 for Rac1). Fold activation was determined relative to the unstimulated controls. (C) Analysis of the Rac effector PAK1/2. T cells were stimulated with S1P or CCL19 for the indicated times, lysed with TritonX- 100 lysis buffer, and analyzed by immunoblotting for phospho-PAK1/2, using GAPDH as a loading control. Top, representative Western Blot. Bottom, quantification from independent experiments (n=3). Fold increases were determined relative to the unstimulated controls. (D-E) Analysis of Rac1 activation in living cells. Purified human CD4^+^ T cells were transduced with Pak-PBD-mCherry, seeded onto fibronectin-coated glasses, and imaged before and after addition of 125nM S1P or 50mM CCL19 for up to 9min. Dark grayscale indicates Rac1-GTP levels. D, representative time-lapse images. E, Quantification of Rac1 activity at the membrane. Rac1-based protrusions were recorded, and their fluorescence intensity was compared to that of the same membrane area at steady state (dotted line). Plotted is the average Rac1 activity for S1P (blue) and CCL19 (red) over time. n = at least 20 separate cells for each condition, obtained from two separate experiments. For all panels, data represent means +/- StDev from multiple experiments. Statistical analysis was performed using an unpaired Student’s T test, relative to the unstimulated control. *, p<0.05; **, p<0.01; ***, p<0.005.

RhoA activation was detectable within 30 seconds of stimulation, and was sustained for at least 5 minutes. In contrast, CCL19-stimulated cells did not exhibit detectable RhoA activation, but these cells showed elevated Rac1-GTP levels (Fig. 2A and B, dark bars). In S1P-stimulated cells, low level activation of Rac1 was sometimes detectable using this assay, but this was not consistently observed and the changes did not reach statistical significance in this population-based assay (Fig. 2B, light bars).

Because pull-down assays sometimes lack the sensitivity to detect weak or transient Rho-GTPase responses, we tested the ability of S1P to induce Rac1 activation in two additional ways. First, we analyzed the phosphorylation of the Rac1 effector PAK. As expected, strong and sustained PAK activation was observed in response to CCL19 (Fig. 2C). Some PAK phosphorylation was detectable in S1P stimulated cells at the 30 second time point, but this response was weak and rapidly extinguished. To further explore this finding, we assessed Rac1 activation kinetics by imaging membrane recruitment of a PAK-based biosensor in living cells.

This approach revealed that stimulation with CCL19 induced strong, sustained Rac1 activation at the membrane, whereas stimulation with S1P induced a transient Rac1 response, lasting only about 3 minutes after the addition of chemoattractant (Fig. 2D and E). Thus, we conclude that S1P and CCL19 activate distinct Rho-GTPase responses; CCL19 drives a sustained Rac1 response while S1P elicits only weak, transient Rac1 activation followed by sustained RhoA signaling.

### S1PR1 and Gαi control chemotactic signaling

In addition to S1PR1, naïve T cells express S1PR4 (*10*), which can signal through Gα12/13 to activate RhoA . We previously showed that S1PR1 inhibition impairs S1P- dependent T cell migration (*13*), but this does not rule out a role for S1PR4. To determine which S1P receptor(s) are involved, T cells were pre-treated with the S1PR1 inhibitor Ex26 or the S1PR4 inhibitor CYM50358 prior to stimulation with S1P, and signaling was analyzed by Western blotting and GTPase pulldown. S1P-induced phosphorylation of AKT and ERK was impaired by Ex26, but not CYM50358 (Fig. 3A-C). Similarly, treatment with Ex26, but not CYM50358, blocked S1P-dependent RhoA activation (Fig. 3D-E). Lastly, we assessed the effects of the two inhibitors on T cell migration in transwell assays. Ex26 impaired S1P- dependent migration, while CYM50538 had no effect (Fig. 3F-G).

**Figure 3:**
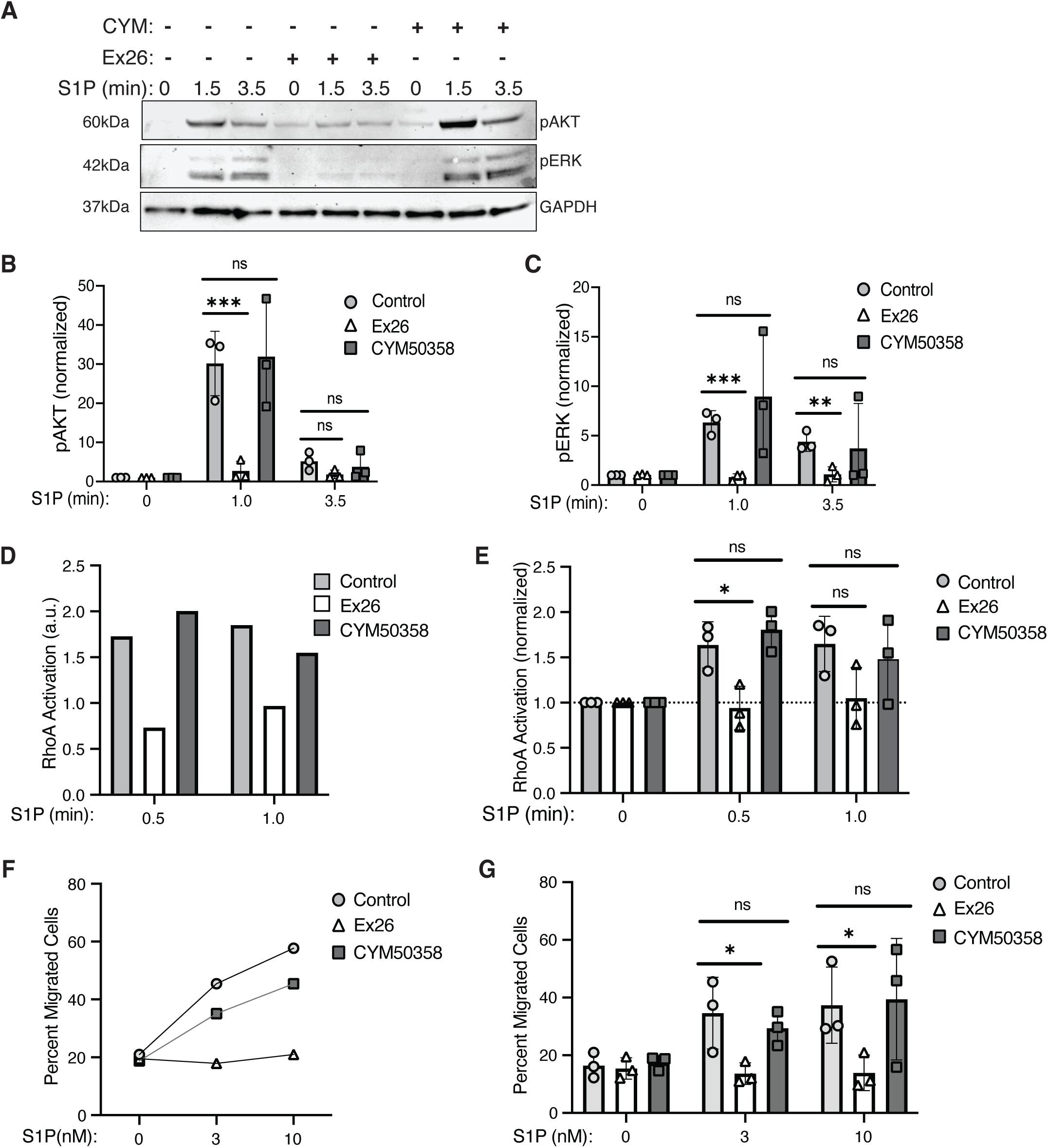
S1P directs naïve T cell signaling and migration via S1PR1. (A-C) Receptor dependence of Akt and Erk signaling. Naïve CD4^+^ T cells were left untreated or pretreated with Ex26 or CYM50358 (CYM) to inhibit S1PR1 or SIPR4, respectively, prior to stimulation with S1P. Lysates were immunoblotted for pAkt, pErk, and GAPDH. (B-C) Quantification of multiple experiments (n=3) performed as in panel A. Fold increases were determined relative to the unstimulated control. (D-E) Receptor dependence of RhoA activation. T cells were pretreated with inhibitors prior to stimulation with S1P, and lysates were analyzed for RhoA-GTP using G-LISA assays. (D) One representative experiment. (E) Quantification of multiple biological replicates (n=3), normalized to the unstimulated controls. (F-G) S1PR1, but not S1PR4, controls S1P-dependent migration. T cells were pretreated with inhibitors, and chemotactic responses were analyzed using transwell assays in the continued presence of inhibitor. (F) One representative experiment, showing a dose-response to S1P. (G) Quantification of multiple biological replicates (n=3), normalized to the unstimulated controls. For panels B,C,E, and G, data represent means +/- StDev from at least three independent experiments. Statistical analysis was performed using an unpaired Student’s T test, relative to the unstimulated control. *, p<0.05; **, p<0.01; ***, p<0.005.

S1PR1 is canonically coupled to Gai (*31, 34, 35*), which usually activates Rac1. However, in our system, engagement of S1PR1 leads to RhoA activation. Gai-dependent activation of the RhoA pathway has been observed downstream of CCR7 in dendritic cells (*36*), but this has not been reported in T cells. To ask if S1P responses in naïve T cells require Gαi signaling, T cells were treated with pertussis toxin (PTX) prior to analysis. PTX treatment inhibited S1P-dependent phosphorylation of Akt and Erk (Fig 4A-C), as well as activation of RhoA (Fig 4D,E). Migration toward S1P in transwell assays was also PTX sensitive (Fig 4F,G). Taken together, these results show that S1PR1, but not S1PR4, is responsible for S1P- dependent signaling and chemotaxis in naïve T cells, and that it does so by coupling to Gai.

**Figure 4:**
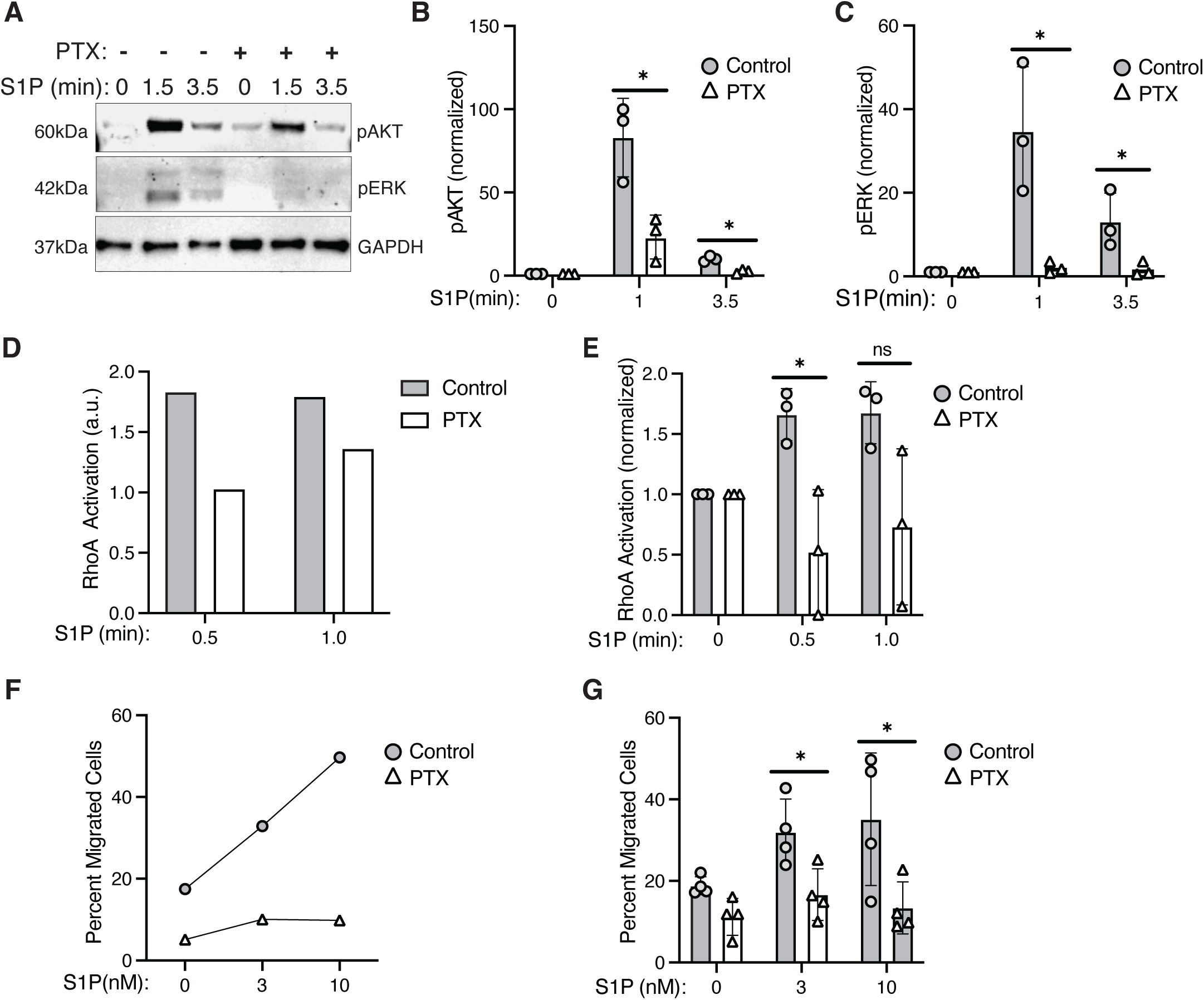
S1PR1 responses require Gai. (A-C) Pertussis toxin inhibits S1P-dependent phosphorylation of Akt and Erk. A, Naïve CD4+ T cells were left untreated or pretreated with pertussis toxin (PTX) to inhibit Gai, prior to stimulation with S1P. Lysates were immunoblotted for pAkt, pErk, and GAPDH. (B-C) Quantification of experiments performed as in panel A. Fold increases were determined relative to the unstimulated control (n=3 independent experiments). (D-E) PTX inhibits S1P-dependent RhoA activation. T cells were pretreated with PTX prior to stimulation with S1P, and lysates were analyzed for RhoA-GTP using G-LISA assays. (D) One representative experiment. (E) Quantification of multiple biological replicates (n=3), normalized to the unstimulated controls. (F-G) PTX inhibits S1P-dependent migration. T cells were pretreated with PTX and chemotactic responses were analyzed using transwell assays in the continued presence of inhibitor. (F) One representative experiment, showing a dose-response to S1P. (G) Quantification of multiple biological replicates (n=4), normalized to the unstimulated controls. For panels B,C,E, and G, data represent means +/- StDev from multiple experiments. Statistical analysis was performed using an unpaired Student’s T test, relative to the unstimulated control. *, p<0.05.

### Receptor-proximal signals are linked to Rho-pathway activation by mTORC2

We next sought to understand how Gai signaling leads to activation of Rho. Recent work from De Belly et al. (*37*) revealed that generation of Rac-based actin-rich protrusions using either optogenetic approaches increases membrane tension, which is sensed by mTORC2 to yield RhoA activation at the opposite pole of the cell. Since we observed that S1P-treated T cells undergo transient Rac1 activation (Fig. 2), and since these cells exhibit a period of membrane ruffling prior to the onset of bleb formation (*13*), we wondered whether an analogous relay system is playing a role in S1P responses (Fig. 5A). To test this, we treated T cells with the mTORC2 inhibitor KU-0063794 and tested S1P-dependent signals leading to bleb formation. As hypothesized, we found that mTORC2 inhibition blocked S1P-dependent phosphorylation of MLC (Fig. 5B,C). This points to a model in which initial S1P signals induce transient Rac-1 activation, leading to the formation of actin-rich ruffles and lamellipodial protrusions. The formation of these protrusions then elevates membrane tension and activates mTORC2, which in turn activates Rho-dependent pathways leading to bleb-based motility.

**Figure 5:**
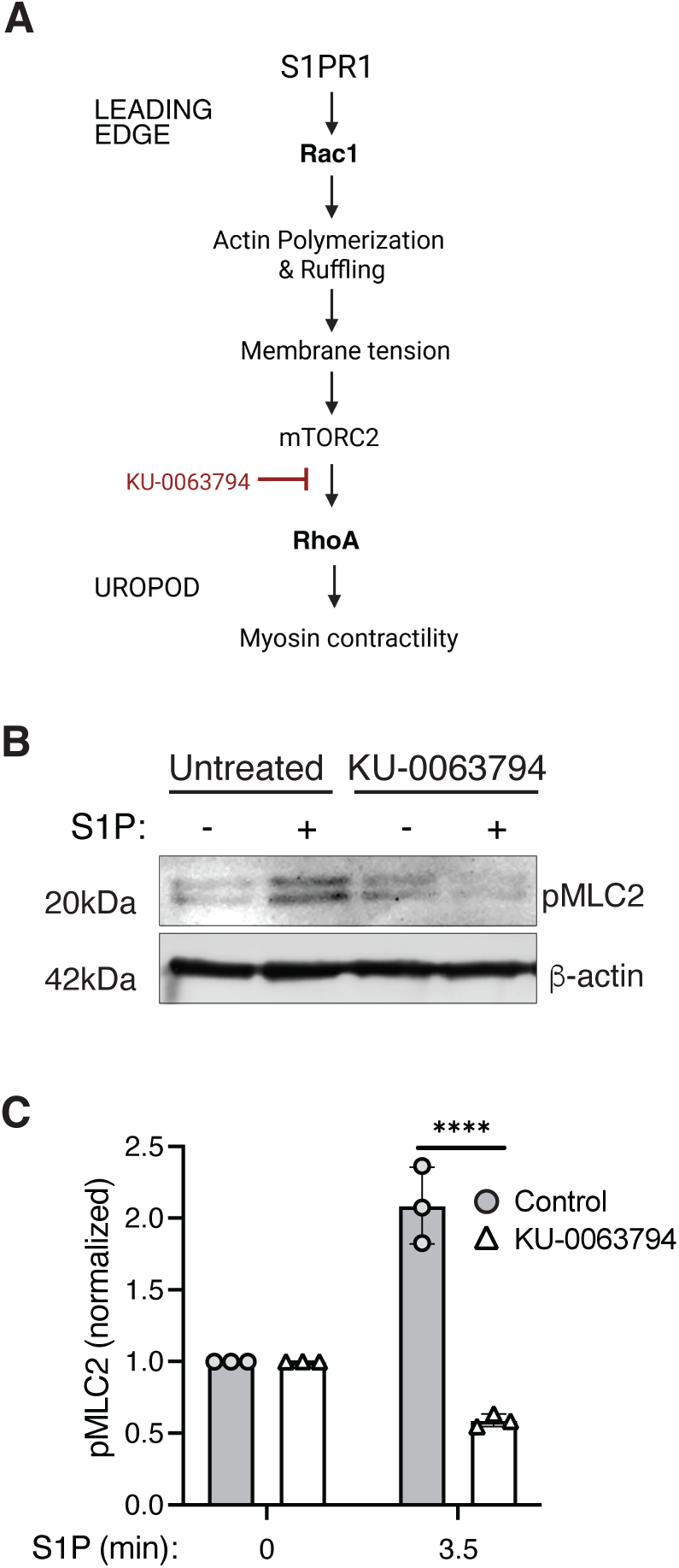
mTORC2 activates RhoA. (A) Proposed model through which initial Rac1-dependent protrusions lead to RhoA activation, with inhibitor indicated in magenta. Early S1P-dependent Rac1 signaling induces actin polymerization and ruffling. This creates increased membrane tension, which is sensed by mTORC2 to activate RhoA. RhoA signaling drives actomyosin contractility at the rear of the cell. (B-C) pMLC increase occurs downstream mTORC. T cells were incubated for 1.5hr with the mTORC inhibitor KU-0063794 or control media prior to stimulation with S1P. Cells were lysed with hot SDS and samples were immunoblotted for pMLC and β-actin. (B) Representative Western Blot. (C) Quantification from independent experiments (n=3). Fold increases were determined relative to the unstimulated controls. Data represent means +/- StDev from multiple experiments. Statistical analysis was performed using an unpaired Student’s T test, relative to the untreated control. ****, p<0.001.

### S1P activates WNK kinases, which function together with myosin to promote migration

Our data show that S1P signaling through the RhoA pathway leads to enhanced MLC phosphorylation, and we showed previously that inhibiting myosin contractility blocks S1P- induced bleb formation (*13*). We therefore expected that inhibiting myosin contractility would block S1P-induced naïve T cell migration. Surprisingly, however, we found that inhibitors of ROCK (Y27632), myosin light chain kinase (ML-7), and myosin ATPase activity (Blebbistatin), all of which inhibit blebbing (*13*), failed to inhibit S1P-induced chemotaxis in transwell assays (Fig 6A). Thus, inhibition of myosin-induced blebbing is not sufficient to block migration; other processes must play a role.

**Figure 6:**
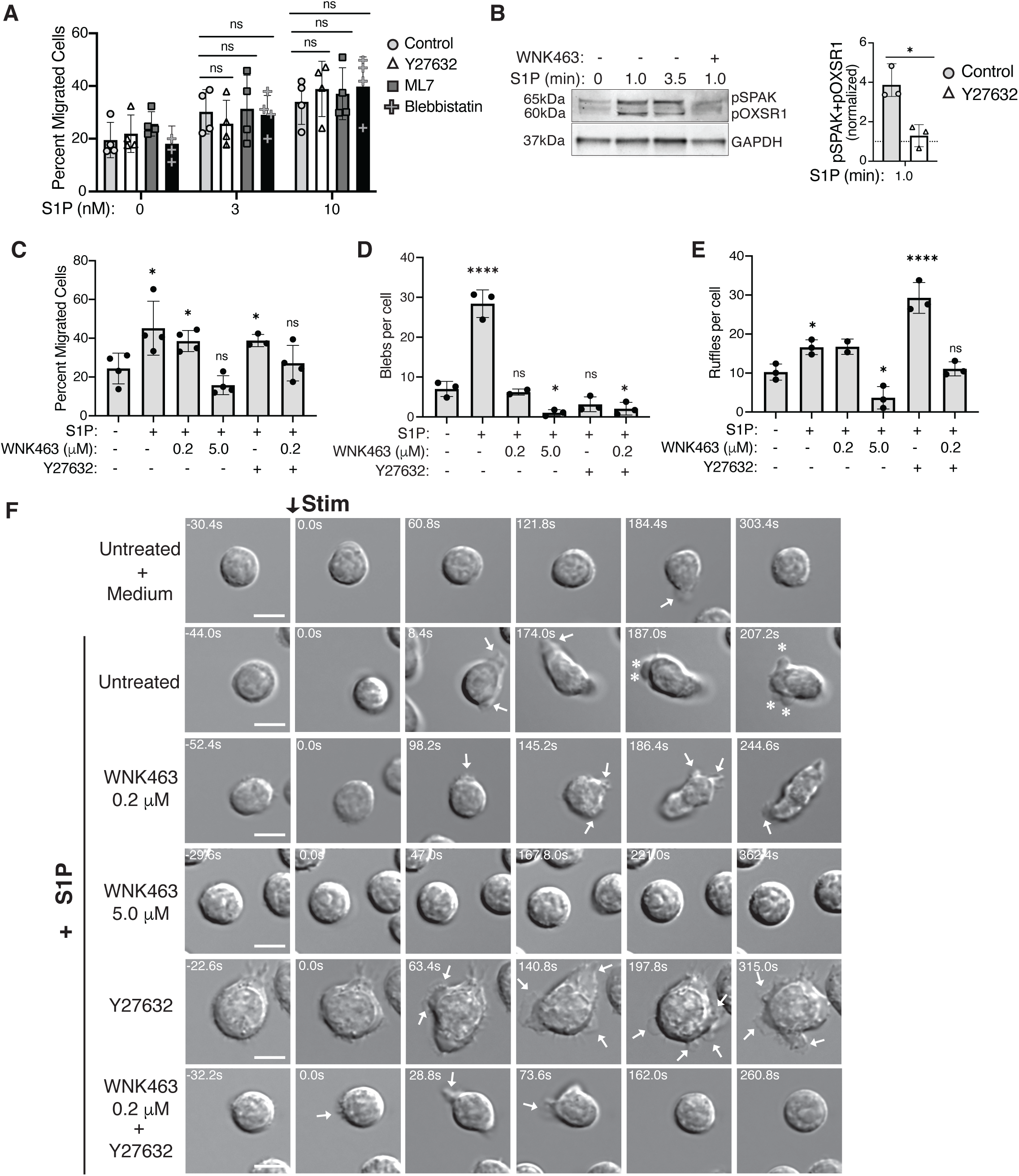
S1P activates WNK kinases, which function together with myosin to promote migration. (A) T cells do not require myosin contractility to migrate toward S1P in transwell assays. T cells were pre-treated with the myosin pathway inhibitors Y-27632, ML-7, or blebbistatin, or with control media. Migration toward the indicated doses of S1P was measured using transwell assays, in the continued presence of inhibitor. n=4 independent experiments. (B) S1P activates WNK1-dependent phosphorylation of ion channel kinases. T cells were incubated for 1.5hrs with the WNK-1 inhibitor WNK463 (5μM) or with control media prior to stimulation with 100nM S1P. Lysates were immunoblotted for phospho-SPAK / phospho-OXSR1 and GAPDH. Left, one representative Western Blot. Right, quantification from independent experiments (n=3). Fold comparisons were drawn to the unstimulated control (dotted line). (C) T cell migratory responses after inhibition of WNK1 and myosin signaling. T cells were left untreated, or pre-treated with 0.2 or 5μM WNK463, 25μM Y-27632, or 0.2μM WNK463 together with 25μM Y-27632. Chemotaxis toward 10nM S1P was tested using transwell assays, in the continued presence of inhibitor(s). n=4 independent experiments. (D-F) Video analysis of T cell protrusions after inhibition of WNK1 and myosin signaling. T cells were pretreated as in panel C, and adhered to VCAM-1 coated coverslips. Cells were imaged using DIC optics before and after the addition of 100nM S1P or control media. Video sequences from independent experiments were manually analyzed for the frequency of ruffles (D) and blebs (E) after stimulus addition as described in Materials and Methods. n=3 independent experiments. (F) Timelapse sequences from representative T cell motility videos. Time is in seconds relative to the point of stimulation. Arrows indicate protrusions classified as ruffles; asterisks indicate protrusions classified as blebs. Bars = 5μm. Data from all panels represent means +/- StDev from multiple experiments. Statistical analysis for panels A-C was performed using an unpaired Student’s T test, relative to the untreated (A,B) or unstimulated (C) controls. Statistical analysis for panels D-E was performed using a one-way ANOVA, relative to the media control, with Dunnett’s multiple comparison’s test used to derive p values. *, p<0.05; ****, p<0.001.

Recent work from the Tybulewicz lab showed that WNK kinases promote CCL21- dependent T cell motility (*38, 39*). WNK kinases phosphorylate OXSR1 and SPAK/STK39, which in turn phosphorylate and activate the ion channel SLC12A2, driving aquaporin-dependent water entry. In T cells responding to CCL21, this pathway fosters motility by releasing membrane-cortex attachment prior to the generation of actin-rich protrusions (*39*). Release of membrane-cortex attachment is also a prerequisite for the formation of bleb-based protrusions in other systems (*40–42*), and we showed that it plays a crucial role in S1P- stimulated naïve T cells (*13*). In addition to its effects on membrane-cortex attachment, water entry elevates intracellular pressure (*43, 44*), and could help generate the forces that drive bleb formation (*24, 25*). Based on this rationale, we asked if stimulation with S1P activates the WNK- signaling cascade. As shown in Fig. 6B, stimulation with S1P resulted in phosphorylation of OXSR1 and SPAK/STK39, and this process was inhibited by pretreatment with the pan-WNK inhibitor WNK463 at 5μM (the same dose shown to inhibit responses to CCL21 (*39*)). 5μM WNK463 also fully blocked chemotaxis toward S1P in transwell assays (Fig. 6C). To ask if the WNK response works in concert with myosin contractility, we treated cells with a reduced dose of the drug (0.2μM), which was previously shown to reduce speed, but not block motility, in CCL21-stimulated cells (*39*). At this dose, WNK463 did not diminish S1P-dependent transmigration. However, when low-dose WNK463 was combined with the myosin pathway inhibitor Y27632 (which also does not block motility on its own), S1P-dependent T cell migration was effectively blocked. These results show that WNK-dependent responses work together with myosin contractility to promote S1P-dependent T cell migration.

### Video analysis of S1P-induced protrusions

To better understand how WNK and myosin influence migration, we examined motile responses using video microscopy. T cells were treated with inhibitors or left untreated, loosely adhered to VCAM-1 coated surfaces, and imaged by DIC microscopy before and after the addition of S1P. Time-lapse movies were analyzed for the frequency of bleb (Fig. 6D) or ruffle (Fig. 6E) formation. Representative time-lapse images are shown in Fig. 6F and the corresponding Supplemental Movies 1-6. As anticipated, analysis of the blebbing response revealed that control cells stimulated with media alone blebbed infrequently, and that the addition of S1P induced robust blebbing (Fig. 6D and F; Movies S1 and S2, asterisks).

Consistent with the finding that complete inhibition of WNK kinase blocks membrane-cortex detachment (*40*), treatment with 5µM WNK463 prior to S1P stimulation blocked almost all movement (Movie S4). The S1P-induced blebbing response was inhibited by treatment low dose WNK463 (Movie S3) or with Y27632 (Movie S5), indicating that both water influx and myosin contractility contribute to the blebbing response. Consistent with this, the blebbing response was further reduced in cells treated with these two drugs in combination (Movie S6). We next analyzed membrane ruffle formation in the same set of movies. Consistent with our data showing transient Rac1 activation, we found that treatment with S1P increased the frequency of ruffle formation (Fig. 6E and F; Movies S1 and S2, arrows). This ruffling response dominated at early times after stimulation and continued throughout the observation period, even after blebbing frequency increased. As with bleb formation, ruffling activity was abrogated by treatment with high-dose WNK463. In contrast, low-dose WNK463, which was sufficient to significantly perturb blebbing, had little effect on ruffling. Interestingly, we found that treatment with Y27632 dramatically enhanced ruffling activity. However, when Y27632 was administered together with low-dose WNK463, ruffling was restored to baseline levels. Taken together with the corresponding transwell data (Fig. 6C), this analysis of protrusive behaviors indicates that the continued ability of cells treated with Y27632 to migrate in transwell assays is attributable to their increased ruffling activity. In support of this, the combination of Y27632 and low-dose WNK463 blocked both enhanced ruffling and transmigration. More generally, these inhibitor studies demonstrate that S1P-mediated motility involves the coordinated activation of pathways leading to both water influx and myosin contractility. These pathways work together to generate the protrusive forces that drive cell short-range chemotaxis.

## Discussion

Our work provides a comprehensive analysis of the chemotactic signaling mechanisms through which S1P directs bleb-based motility of naive T cells. Our findings reveal a complex mechanism that regulates cell motility through the coordinate function of Rho-GTPases, acto- myosin dynamics, and osmoregulation (modeled in Figure 7). We show that S1P engages S1PR1, which signals through Gai to initiate two complementary signaling cascades, one involving Rac1 and PLC, and another involving WNK1. Activation of Rac1 induces transient actin polymerization followed by sustained RhoA activation. This, together with Ca^2+^ responses induced by PLC signaling, leads to enhanced myosin contractility. In parallel with this, S1P activates a WNK1 signaling pathway resulting in water influx. Myosin contractility and water influx work together to elevate intracellular pressure, driving bleb formation and forward cell movement.

**Figure 7:**
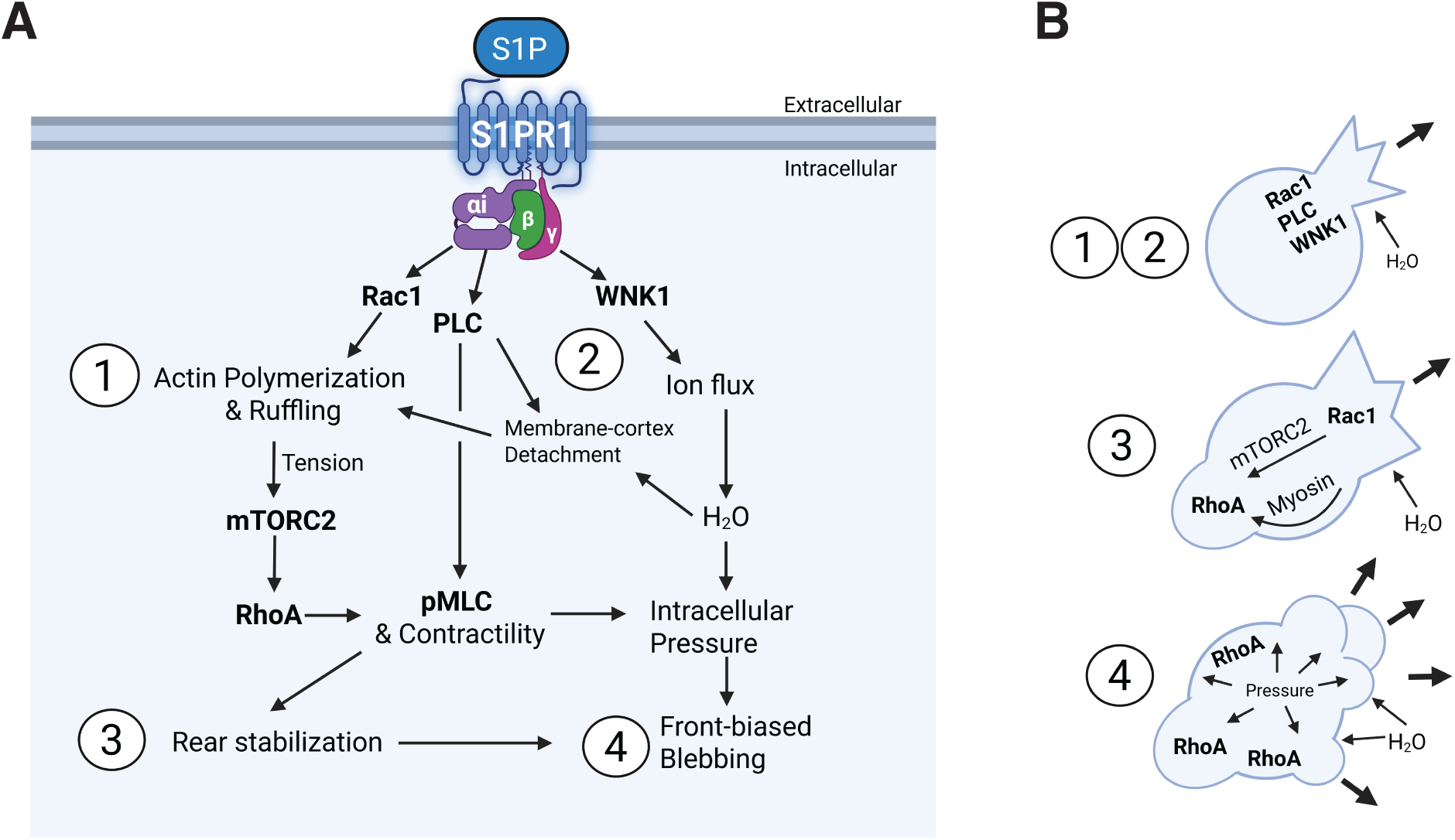
Proposed mechanism through which S1P elicits bleb-based motility of naïve T cells. (A) Key signaling events and (B) associated cell shape changes. Step 1, Naïve T cells bind S1P via receptor S1PR1, and initiate signaling through Gai. This activates Rac1 transiently, which leads to branched-actin filament nucleation, ruffling and lamellipodia. Step 2) Simultaneously with Step 1, S1PR1/Gai activates WNK1-dependent ion-channel phosphorylation, leading to an influx of water from the extracellular space. Water influx works together with PLC-dependent cleavage of PIP2 to favor detachment of the plasma membrane from the actomyosin cortex at the leading edge. This allows Rac1 activity to form actin-rich membrane ruffles, thereby generating membrane tension. Step 3) Membrane tension is sensed by mTORC2, which then activates RhoA. RhoA, along with Ca^2+^ signaling events downstream of PLC, activates actomyosin contractility. The resulting rearward flow and concomitant cortical reattachment forms a stable uropod in the rear of the cell, reinforcing cell polarity and biasing protrusions to the leading edge. Step 4) RhoA-dependent actomyosin contractility also increases intracellular pressure, as does WNK1-dependent water influx. This pressurization generates transient blebs at the front of the cell, where membrane-cortex attachment is weakest. Blebs are then retracted by actin filaments, allowing the cell to form new actin- or fluid-driven protrusions. According to this model, initial membrane-cortex detachment and transient Rac activation are essential for establishing cell polarity and initiating motility, while ongoing actin polymerization and/or elevated intracellular pressure drive the formation of leading-edge protrusions that power forward cell migration.

To our knowledge, this is the first report showing that S1P activates WNK1 signaling in T cells. We show that S1P induces WNK-dependent phosphorylation of OXSR1/SPAK kinases, activating a pathway known to induce water influx and elevate intracellular pressure. This pathway works in conjunction with myosin contractility to drive S1P-dependent chemotaxis, as simultaneous inhibition of both pathways blocked chemotaxis under conditions where inhibition of the individual pathways did not. The WNK pathway is also activated in CCL21 stimulated T cells, where it was proposed to promote release of membrane-cortex linkers (*39*). In keeping with this, we found that complete WNK inhibition blocked chemotactic responses to S1P. Indeed, this treatment arrested nearly all cell motility, consistent with the idea that release of membrane-cortex attachment is an essential first step for formation of all protrusions, including those driven by intracellular pressure (*40–42*). Thus, we propose that WNK pathway activation plays a dual role in the S1P response, regulating membrane-cortex attachment and working in tandem with myosin responses to increase intracellular pressure and drive bleb-based motility.

Our results shed important new light on how CCL19 and S1P induce distinct modes of T cell motility. As previously shown by others (reviewed in (*7, 45, 46*)), stimulation of T cells with CCL19 induces sustained Rac1 activation, leading to branched actin polymerization and sustained motility driven primarily by lamellipodial protrusions. In contrast, we find that S1P induces very transient Rac1 activation, followed by a sustained RhoAresponse that orchestrates short range, directional, bleb-based motility. The initial, transient Rac1 response observed in S1P-stimulated T cells induces ruffling similar to that induced by CCL19. Along with localized PLC activity, this transient burst of actin polymerization likely establishes cell polarity by defining the site where membrane-cortex attachments are weak and protrusive structures can form. This early Rac1 activation then transitions into a dominant RhoAresponse. The resulting increase in myosin contractility (in conjunction with water influx) elevates intracellular pressure and favors bleb-based protrusive forces.

One surprising finding in this study was the activation of RhoA downstream of S1PR1 and Gai. Multiple S1PRs are expressed in different T cell subsets (*47*), some of which are known to activate RhoAthrough coupling to Ga12/13. These include S1PR4, which is expressed along with S1PR1 in naïve T cells, and which has been implicated along with S1PR1 in S1P- driven transendothelial migration (*10*). Using receptor-specific antagonists, we found that inhibition of S1PR1 was sufficient to fully block both signaling and transmigration, whereas inhibition of S1PR4 had little or no effect. In addition, PTX completely blocked the S1P response, consistent with a pathway involving S1PR1 and Gai. Insights into how S1PR1/Gai could activate RhoA come from work by De Belly et al. (*37*), who showed using CCL19-treated cells and optogenetic approaches that RhoA can be activated downstream of Rac1. In that system, Rac1-mediated actin polymerization at the leading edge creates membrane tension, which is sensed by mTORC2. This, in turn, activates RhoA and actomyosin contractility at the rear of the migrating cell. This mechanism also appears to be at play in S1P-stimulated T cells, since we find that mTORC activity is required for S1P-dependent MLC phosphorylation.

Moreover, our live-cell imaging shows that treatment with drugs that prevent membrane ruffling also blocked bleb formation, whereas the converse was not true. Thus, our results are consistent with a model in which S1P signals establish a 2-phase actin-regulatory response whereby an initial period of Rac-dependent actin polymerization sets the stage for a Rho-dependent phase of increased actomyosin contractility. One appealing aspect of this model is that it explains how the polarity of bleb-based motility is established, since the initial phase of polarized actin polymerization defines the leading edge of the migrating cell.

One of the most striking differences in the responses to CCL19 vs S1P is in the kinetics of Rac1 activation. Using a live-cell Rac1 reporter, we found that Rac1 activation is sustained for over 10 minutes in cells stimulated with CCL19, but extinguished within about 3 minutes of S1P stimulation. This finding is consistent with our earlier working showing that S1P induces more transient increases in F-actin content, and shorter duration of cell motility (5-7 minutes for S1P vs >15 minutes for CCL19 (*13*)). What controls the brief duration of Rac1 signaling in S1P- treated cells is currently unknown. One possibility is short-range inhibition by Rho, although this is somewhat difficult to reconcile with our observation that mTORC2 signaling is needed to activate Rho. A more appealing possibility is that the duration of Rac1 signaling is dictated by the rates of receptor internalization and signal extinction. S1PR1 is very rapidly internalized, leading to signal termination (*12, 48*). In contrast, several processes can delay CCR7 signal termination: receptor internalization can be delayed by glycosylation, the receptor can recycle back to the surface, and it can signal to Rac1 from intracellular pools (*49–51*). These differences in receptor trafficking could differentially regulate Rac1 GEFs or the inositol lipids that regulate their activity, such that CCL19 signals yield a sustained response, and S1P a shorter one. Going forward, this will be an important area for future investigation.

Several recent studies show that release of membrane-cortex attachment is required for generation of both actin-rich and bleb-based protrusions (*40–42*). Thus, it will also be important to understand the mechanisms that coordinately regulate cortical attachment and protrusive force. This is especially important for S1P, since we have shown that polarized bleb formation and chemotaxis depend on the ERM-family cortical linker protein moesin (*13*). Both arms of the S1P signaling pathway that we have defined are expected to lead to the release of membrane-cortex attachment. By cleaving PIP2, PLC depletes binding sites for ERM proteins, initiating their rapid release from the plasma membrane (*52*). Similarly, WNK1-dependent water influx has been shown to drive separation of the membrane from the underlying actin cortex, and to sustain depletion of membrane-cortex attachment sites at the leading edge of migrating cells (*40*). In support of the idea that both PLC and WNK1 mediate release of the actin cortex, treatment of T cells with inhibitors of either PLC or WNK1 result in rounding up and near-complete loss of motility ((*39, 52*) and Figure 6F).

In addition to this initial cortical release mechanism, membrane-cortex attachment and protrusive force appear to be coupled through feedback mechanisms involving Rho-GTPases. We showed previously that S1P induces highly transient ERM protein dephosphorylation (2-4 minutes), whereas CCL19 induces much more sustained ERM protein dephosphorylation (*13*). These kinetics correlate well with the kinetics of Rac1 activation in the current study, but the causal relationships remain unclear. Differential effects on inositol lipids can explain both sets of events, since activation of ERM proteins is initiated by binding to plasma membrane pools of PIP2 (*53*). However, the two sets of responses may be linked in other ways as well. Rac1 activation has been long been known to correlate with ERM protein dephosphorylation (*54–56*), whereas RhoA activation fosters ERM phosphorylation. RhoA-GTP binds to the ERM kinases SLK and LOK (*57*), and ROCK inhibits MLCP, the phosphatase that works on both myosin and moesin (*22, 58*). Rho-GTPases and ERM proteins are also interlinked via subcellular localization. When ERM proteins are rapidly re-phosphorylated in S1P-stimulated cells, they are swept toward the back of the cell along with the contractile actomyosin network. This effectively reinforces the back of the cell and weakens the front, allowing pressure-induced forces to generate a polarized motile response.

From an immunological perspective, it will be important to ask how T cells utilize their migratory versatility to sense environmental cues and move through different types of tissue barriers. Lamellipodial motility allows cells to traverse long distances along adhesive surfaces, especially in relatively unconfined tissue settings. In contrast, bleb-based movement can occur in the absence of adhesive ligands, provided that the cell is at least partially confined. Indeed, bleb-based motility allows cells to rapidly sense and find a path through narrow openings within dense tissue stroma (*59–61*). This mode of motility also favors frequent turning events, like those needed for efficient antigen search (*62, 63*). Importantly, the Rac1- and RhoA-based motile responses described here probably rarely occur independently of one another. *In vivo,* most T cell responses would involve switching between these two modes, depending on environment cues including chemoattractants, adhesive ligands, and spatial confinement.

Different T cell populations likely exhibit distinct motile behaviors, due to cell-intrinsic set- points for actin regulatory pathways. The studies described here were performed on naïve T cells, which have been previously shown to have higher levels of RhoA and ROCK activity than effector cells (*64*). This was proposed to make naïve cells stiffer and less prone to activation than effectors, but it might also bias them toward bleb-based motility. On the other hand, Ruef et al. (*60*) showed in a salivary gland model that Trm cells, but not naïve T cells, engage in bleb-based motility, and that this depends on confinement-induced activation of myosin contractility. Different effector populations may also have different programmed behaviors. For example, in a model of inflamed skin, Th2 cells turn more frequently and survey a larger area than Th1 cells (*65*). This was shown to be an integrin-dependent response, but that does not preclude a role for differential Rho-GTPase signaling. To fully understand T cell trafficking during an immune response, it will be necessary to determine how signals from multiple chemoattractants, confinement conditions and adhesive interactions converge on developmentally biased actin-regulatory pathways to shape the behavior of specific T cell populations.

## Materials and Methods

### T cell purification and culture

C57BL/6J mice were purchased from Jackson Laboratories and housed in the Children’s Hospital of Philadelphia animal facility, in accordance with protocols approved by the Institutional Animal Care and Use Committee. Mice (both male and female) were sacrificed as a source of T cells at 8-11 weeks of age. Lymph nodes and spleen were harvested and single cell suspensions were prepared in cold MACS buffer (0.5% Fraction V lipid-free BSA (Roche), 1mM EDTA, in PBS). Naïve CD4+ T cells were isolated by negative selection using a magnetic column (Miltenyi Biotec). Cells were sensitized to chemoattractant signaling as described previously (*13*). Briefly, cells were washed four times with MACS buffer, resuspended at 1x10^6^ cells/ml in DMEM (Corning #10-017-CV) with β-mercaptoethanol, GlutaMAX, MEM-Non-essential amino acids, sodium pyruvate, and penicillin-streptomycin (T cell media), supplemented with 10% charcoal-stripped FBS (CS-FBS; Gibco), and then cultured overnight at 37°C and 10% CO_2_ prior to use.

### Chemoattractants

S1P d18:1 powder (Avanti Polar Lipids) was solubilized in methanol:water 95:5 at 50°C with sonication. The solvent was evaporated using dry nitrogen to create a film of S1P on the interior of the vessel; S1P was then resuspended in Milli-Q water supplemented with 4 mg/ml of fatty acid–free BSA (Roche) to yield a 100 µM stock solution. The S1P-BSA stock solution was stored at −20°C in tightly sealed glass vials and diluted into T cell media immediately before use. Recombinant carrier-free murine CCL19 (R&D Systems) was reconstituted at 100μg/ml in PBS, aliquoted, and stored at -80°C for up to three months. Stock solutions were diluted into cell culture media immediately before use.

### Antibodies

The following primary antibodies were used: mouse anti-GAPDH (Millipore Sigma; #MAB374), mouse anti-β-actin (Millipore Sigma, # A5441), rabbit anti-phospho-SPAK (Ser373) / phospho-OSR1 (Ser325) (Millipore Sigma, #07-2273), rabbit anti-phospho-Erk1/2 (Thr202/Tyr204) (Cell Signaling; #9101), rabbit anti-phospho-Akt (Ser473) (Cell Signaling; #4060), rabbit anti-phospho-PAK1 (Ser199/204)/PAK2 (Ser192/197) (Cell Signaling, #2605), rabbit anti–phospho-myosin light chain (Thr18/Ser19) (Cell Signaling; #3674).

### Inhibitors

Inhibitors were obtained from the following sources, and diluted in T cell media to the indicated final concentrations: Blebbistatin (5μM, Cayman Chemical); CYM50358 (25nM, Tocris Bioscience); Ex26 (100nM, Tocris Bioscience); ML-7 (10μM, Calbiochem); U73122 (100nM, Calbiochem); Y-27632 (25μM, Calbiochem); XMU-MP-1 (70nM, Selleck Chemicals); WNK463 (0.2μM or 5μM, Selleck Chemicals); KU-0063794 (10μM, Selleck Chemicals); PTX (200ng/ml, Tocris). For most inhibitors, cells were pre-treated for 20 minutes and maintained in the culture throughout the experiment unless otherwise indicated. 1hr pre-incubation was used for XMU-MP-1 and KU-0063794, while 1.5hr pre-incubation was used for PTX.

### Transwell migration assays

After chemoattractant sensitization, T cells were resuspended in DMEM, 10% CS-FBS containing 2 drops per ml of NucBlue stain (Thermo Fisher). Cells were adjusted to 1x10^6^/ml, and 100 μl (1x10^5^ cells) per well was added to 24-well transwell inserts (polycarbonate 5μm pore, Corning). Cells were allowed to settle for 20 min at 37°C. For inhibitor studies, drugs were premixed with cells prior to adding to the top chamber, continuing pre-treatment during the settling period. The insert was gently lowered into a well containing 600μl media in the presence or absence of chemoattractant (and inhibitor, where applicable). After 90 min, transwell inserts were gently removed and the cells in the lower chamber were resuspended and counted using a Cytoflex 6 flow cytometer after gating on NucBlue-positive cells. A fixed volume of cells was counted; an input control well was used to derive migration percentages.

### Immunoblot analysis

For biochemical analysis of chemoattractant responses, cells were adjusted to 5x10^6^/ml in T cell media containing 10% CS-FBS; 200 μl (1x10^6^ cells) was added to an Eppendorf tube for each time point, and rested for 20min at 37°C. Cells were then stimulated by the addition of 50 μl of 5× S1P (100nM final concentration) or 5x CCL19 (100ng/ml final concentration). To stop, 1 ml of ice-cold PBS was added, cells were rapidly microfuged at approximately 20,000xg for 20 sec, and cell pellets were resuspended in TX-100 lysis buffer (50 mM Tris-HCl, 50 mM NaCl, 5 mM EDTA, 50 mM NaF, 30 mM Na_4_P_2_O_7_, 50 mM β-glycerophosphate, 1X EDTA-free protease inhibitor cocktail (Roche), and 1% Triton X-100). Cells were lysed on ice for 30 min with occasional vortexing, microfuged at 20,000xg for 10 min to remove nuclei and debris, and supernatants were transferred to a fresh tube. For immunoblots aimed at probing for phospho-MLC, cell pellets were instead resuspended in 95°C SDS lysis buffer (50 mM Tris-HCl, 300 mM NaCl, 1 mM EDTA, 50 mM NaF, 30 mM Na_4_P_2_O_7_, 1X EDTA-free protease inhibitor cocktail, and 1% SDS) and boiled for 8 min with pipette resuspension every minute, after which cells were microfuged and post-nuclear supernatants were transferred to a fresh tube. Irrespective of lysis method, lysates were moved to tubes containing NuPAGE LDS sample buffer (Thermo Fisher Scientific) and 0.2M DTT, and heated to 95°C for 5 min. Samples were then resolved on 10% or 4–12% gradient Bis-Tris gels (NuPAGE), transferred to nitrocellulose membranes, and blocked with Intercept Blocking Buffer (LI-COR) diluted 1:1 in PBS. Membranes were then probed overnight at 4°C with primary antibodies diluted in Tris-buffered saline, 0.1% TWEEN 20 (TBST), with 5% (wt/vol) BSA. Following overnight incubation, membranes were washed multiple times with TBST, stained for 1hr at RT with anti-rabbit or anti-mouse secondary antibodies conjugated to AlexaFluor680 or AlexaFluorPlus800 (Thermo Fisher), and washed again. Immunoblots were scanned on a LI-COR Odyssey imaging system and quantified using ImageStudioLite (v. 5.2.5, LI-COR), taking care to remain within the linear range. Protein mobilities, determined based on molecular weight standards (Biorad, #1610374), are indicated on each blot in kilodaltons.

### Biochemical analysis of Rho-GTPase Activation

Timed stimulations were conducted as described above for immunoblotting, except that cell pellets were lysed in ice-cold 1X G-LISA buffer (Cytoskeleton, Inc.) with 50 mM NaF, 30 mM Na_4_P_2_O_7_, and 1X EDTA-free protease inhibitor cocktail, taking care not to disturb the lysates by excess pipetting. Lysates were snap-frozen in liquid nitrogen and stored for a maximum of 1 week at -80°C. Samples were then thawed slowly on ice before pipetting into G-LISA assay plates coated with antibodies to Rac1 or RhoA (Cytoskeleton, Inc.). Plates were incubated with primary antibodies for 30 min at 4°C, with shaking at 400rpm. As positive controls, lysates from untreated cells were activated with 0.2mM GTPγS and assayed in parallel. G-LISA processing and analysis were carried out per manufacturer’s instructions. Background signal (lysis buffer alone) was subtracted from all samples and fold analysis was done by comparing all experimental samples to the non-stimulated control group. Three or more independent experiments were performed, with 2-3 technical replicates each. Statistical comparisons were performed using an unpaired Student’s T test. Means +/- StDev are shown.

### Biosensor-based analysis of Rac1 activation

Biosensor studies were conducted essentially as previously described (*37*). Briefly, blood specimens from healthy human donors were obtained with informed consent according to the IRB-approved study protocol at the University of California - San Francisco (Study #21-35147). Primary human CD4^+^ T cells were isolated by magnetic bead purification, activated with CD3/28 Dynabeads (Gibco, #11131D), and transduced with recombinant lentivirus expressing Pak-PBD-mCherry. On d6, cells were harvested, beads were removed, and cells were sorted based on mCherry expression. Cells were then rested in the presence of 40 U/mL of human IL-2 (NCI BRB Preclinical Repository) for 2 days. Prior to use, cells were cultured overnight in serum starvation media consisting of Lonza X-VIVO15 (#04-418Q), 0.4% BSA (Sigma-Aldrich, #9048-46-8), 10 mM neutralized N-acetyl L-Cysteine (Sigma-Aldrich, #A9165), 1 mM 2-mercaptoethanol (Gibco, #21985-023), and 40 U/mL IL-2. Cells were then seeded on 96-well #1.5 glass-bottom plates (Azenta Life Sciences) pre-coated with human fibronectin at 1mg/ml (Sigma-Aldrich, #F0895). During imaging, starvation media containing 50mM CCL19 (R&D Systems, #361MI025CF) or 125nM S1P was added into the well after 90 sec. For analysis, average Rac1 activity (mCherry fluorescence intensity) was measured at the cell front by manually identifying cell protrusions using a fluorescence Z-stack together with brightfield imaging. To obtain steady state measurements, the same patch of membrane that formed protrusions later was analyzed prior to the addition of chemoattractant.

When a front could not easily be identified, Rac1 activity was measured at the membrane of the mid-plane of the cell. The analysis only included cells that displayed a cell shape change in response to CCL19 or S1P (protrusion, contraction, blebbing, migration etc.).

### Live cell imaging of cell protrusions

8-well chamber slides with #1.5 coverglasses (Lab Tek II, #155409) were coated overnight at 4°C with 2μg/ml recombinant mouse VCAM-1 (R&D Systems, #643-VM-050) in PBS. Wells were then washed with PBS, taking care to prevent drying, and filled with 100μl of L15 imaging media (L15 media without phenol red, Gibco #21083-027, supplemented with 2 g/L glucose). Naïve murine CD4^+^ T cells were isolated and sensitized as described above, except that sensitization was achieved by culturing in T cell media with CS-FBS for 4-5 hr. For experiments involving inhibitors, cells were pretreated during the final hour of sensitization, and then pelleted and resuspended in L15 imaging media. Cells were adjusted to 1x10^6^ cells/ml, and 200ul of cell suspension was added to each well. Slides were transferred to the microscope chamber and cells were allowed to rest and attach for 20min at 37°C before imaging. Differential interference contrast (DIC) imaging was performed at 37°C using a Zeiss Axio Observer 7, equipped with a 63x oil-immersion objective (NA=1.4) and a Zeiss Axiocam 807 sCMOS camera, collecting 1 image every 200msec. One minute after imaging started, 30μl of 10X S1P (100nM final) or L15 alone was added to the well, and imaging was continued for a total of 10min. Analysis of T cell blebbing and ruffling was performed essentially as previously described (*13*), by generating movies in Fiji and scoring manually during playback. To be scored as a bleb, protrusions had to form rapidly (within around 0.2-1s), and to appear smooth, symmetrical, and clear (devoid of any obvious granules or organelles).

Blebs were counted in forward time to avoid confusion with smooth retractions. Analysis of membrane ruffles was conducted similarly, with ruffles defined as individual wave-like membrane protrusions, with complex irregular profiles and grainy contents. In some cases, multiple waves formed within a single protrusive region; this was counted as a single ruffle. Representative MP4 movies were annotated and prepared for publication using Adobe Express.

### Statistical analysis and graphical models

Statistical analysis was performed using Graphpad Prism Software, with specific tests and sample sizes indicated in figure legends. Models and diagrams were prepared using BioRender Software.

## Acknowledgements

The authors thank Dr. Tanner Robertson and Ms. Christine Wu for foundational studies and technical advice and discussion. We thank Dr. Pamela Schwartzberg for advice and critical reading of the manuscript, and Dr. Guilherme Nader and members of the Burkhardt laboratory for many helpful discussions. We thank Mr. Zachary Zimmerman for help with handling stock solutions of S1P, and Mr. Christopher Davis for expert administrative assistance. This work was supported by R01AI147118 and R21AI1739938 to JKB, R35GM118167 to ODW, and K99GM154115 to HDB. FSR was supported by GT15872 from HHMI.

## Supplementary Materials

### Movie Legends

**Movie S1. Naïve T cells show little motility after stimulation with control media.** Naive CD4+ T cells with no prior inhibitor treatment were stimulated with L15 media as a vehicle control. These cells only infrequently formed protrusions over the course of 9min. Protrusions are identified with arrows for ruffles and asterisks for blebs. The indicated time (s) is relative to stimulation.

**Movie S2. S1P induces naïve T cells to ruffle and bleb.** Naive CD4+ T cells with no prior inhibitor treatment were stimulated with 100nM S1P at t=0 sec. These cells formed many protrusions. Over the first few minutes, protrusions mainly consisted of ruffles and then the cells began forming very frequent leading-edge blebs. Protrusions are identified with arrows for ruffles and asterisks for blebs. The indicated time (s) is relative to stimulation.

**Movie S3. WNK463 low dose inhibits bleb formation.** Naive CD4+ T cells were pre-treated with 0.2μM WNK463 and stimulated with 100nM S1P at t=0 sec. These cells rarely formed blebs but formed ruffles as frequently as control S1P-stimulated cells. Protrusions are identified with arrows for ruffles and asterisks for blebs. The indicated time (s) is relative to stimulation.

**Movie S4. WNK463 high dose inhibits bleb formation.** Naive CD4+ T cells were pre-treated with 5μM WNK463 and stimulated with 100nM S1P at t=0 sec. These cells rounded up and rarely formed protrusions of any kind. Protrusions are identified with arrows for ruffles and asterisks for blebs. The indicated time (s) is relative to stimulation.

**Movie S5. Y-27632 inhibits bleb formation.** Naive CD4+ T cells were pre-treated with 25μM Y-27632 were and stimulated with 100nM S1P at t=0 sec. Y-27632 treatment caused notable morphological changes relative to untreated controls; cells became become flatter and more spread. They formed and retracted many simultaneous ruffles, but rarely formed blebs. Protrusions are identified with arrows for ruffles and asterisks for blebs. The indicated time (s) is relative to stimulation.

**Movie S6. Y-27632 and WNK463 low dose inhibits bleb formation.** Naive CD4+ T cells were pre-treated with 25μM Y-27632 and 0.2μM WNK463 and stimulated with 100nM S1P at t=0 sec. These cells looked morphologically similar to control T cells, and lacked the frequent ruffling responses observed in cells treated with Y-276323 alone. These cells produced many fewer ruffles and blebs than control cells stimulated in the absence of inhibitor, though they remained more active than cells treated with 5μM WNK463. Protrusions are identified with arrows for ruffles and asterisks for blebs. The indicated time (s) is relative to stimulation.

## Notes

### Competing Interest Statement

The authors have declared no competing interest.

